# Concordia: Spatial Domain Detection via Augmented Graphs for Population-Level Spatial Proteomics

**DOI:** 10.64898/2026.04.19.719422

**Authors:** Si Liu, Li Hsu, Wei Sun

**Affiliations:** Biostatistics Program, Public Health Sciences Division, Fred Hutchinson Cancer Center, Seattle, Washington, USA; Department of Biostatistics, University of Washington, Seattle, Washington, USA; Department of Biostatistics, University of North Carolina, Chapel Hill, North Carolina, USA

## Abstract

A key step in analyzing population-level spatial proteomic data is to delineate consistently defined spatial domains across samples. Domain detection is particularly challenging for cancer tissues, which have complex spatial domains with elongated or branching geometries. To address these challenges, we present Concordia, a Graph Neural Network (GNN)-based framework that uses augmented graphs to capture complex spatial domains, and it is designed to analyze thousands of tissues simultaneously to obtain consistently defined domains. Applied to a lung cancer dataset, Concordia uncovers a spatially defined cancer associated fibroblast subset linked to clinical outcomes that cannot be identified using protein expression alone.

## Introduction

Spatial proteomics techniques, such as Imaging Mass Cytometry (IMC)^1^ and PhenoCycler (formerly CODEX)^2^ allow profiling of tens of protein markers in single cell resolution. They have been used to generate large spatial proteomic datasets, including hundreds to thousands of tissue samples.^3–5^ These large datasets enable population-level analyses linking spatially organized tissue microenvironments to clinical outcomes. A tissue microenvironment, often referred to as a spatial domain, is characterized by its cell type composition and spatial organization. For example, one spatial domain may be tumor dominant, while another may represent a tumor-immune mixture at the invasion boundary. However, systematically identifying such complex spatial domains across large patient cohorts remains a challenge.

Given the informativeness of spatial domains, many methods have been developed for spatial domain annotation. Most focus on spatial transcriptomic data.^6–8^ Due to the sparsity of transcriptomic data, cell type calling can be challenging and thus domain detection and cell type calling are often combined into one task.^6, 8^ A few methods integrate spatial multiomic data for domain detection,^9, 10^ and others can handle either spatial transcriptomic or spatial proteomic data.^11–13^ Across all of these approaches, quantitative features (e.g., gene expression or protein abundance) have been used as the inputs. We argue that for spatial proteomics, cell type calling is more trackable than spatial transcriptomics. Protein panels are typically designed around cell type markers and their measurements are far less sparse than transcriptomic data. Therefore, for spatial proteomic data, it is feasible and desirable to annotate domains based on cell type calls. This approach offers two advantages. First, cell types can be called within each batch or each sample, and thus the cell type calls are less sensitive to the technical noise and batch effects. Second, such domain detection based on cell types are easy to transfer across datasets and platforms, as cell types are well defined and readily interpretable. Therefore, in this work, we focus on domain annotation using cell type labels and cell coordinates as input.

Broadly, there are three types of computational approaches for spatial domain detection using cell type labels. The first one is to directly cluster cells based on the cell type composition of each cell’s neighborhood.^2, 14, 15^ The second type uses Latent Dirichlet Allocation (LDA) and models tissues as mixtures of latent topics with spatial smoothness priors^16^ or by incorporating distance-based kernels.^17^ The third employs Graph Neural Network (GNN),^5, 18, 19^ which constructs a cell-cell graph and propagates information across cells in this graph to learn new embedding of each cell. In the existing GNN methods,^5, 19^ cell-cell graphs have been constructed to connect nearby cells. For example, a cell may be connected to all the cells within a certain distance or all k-nearest neighbors. LDA is more flexible than direct clustering methods since it allows multiple topics per cell neighborhood. Compared with LDA-based methods such as Spatial-LDA^16^ and SpatialTopic,^17^ a GNN-based method has the flexibility to learn new embeddings for each cell.^19^ All these methods are designed to capture local and compact domains, but they are less effective to detect elongated and branching structures, such as tumor invasive fronts.

To address these limitations, we propose a GNN-based domain detection framework named Concordia, which operates on an augmented cell–cell graph. In this augmented graph, long-distance edges are added based on neighborhood similarity, enabling detection of elongated or branching patterns commonly observed in tumor tissues. Rather than direct graph partitioning, Concordia extracts learned cell embeddings and applies a two-stage clustering procedure that respects both embedding and physical coordinate distances. Crucially, Concordia operates across the entire dataset simultaneously, enabling the discovery of domains that are consistently defined across all images. This is made feasible through a sparse representation of cell-cell graph, which greatly reduces memory usage and thus makes the analysis scalable to datasets with a large number of cells.

The spatial domains identified by Concordia can be directly used for association analyses between domain abundance and clinical outcomes. Applied to a lung cancer cohort, we used Concordia-identified domains to link with survival times and further revealed context-specific effects, whereby the prognostic impact of certain cell types was evident only within certain spatial domains. These findings demonstrate that these spatial domains capture biologically meaningful and potentially novel cell subsets with clinical relevance.

## Results

### Brief introduction to the Concordia method

The Concordia method operates on spatial single-cell datasets in which each cell has a spatial coordinate and a cell-type label. We focus on spatial proteomic data because they provide reliable cell type information and are scalable to larger sample size. Spatial transcriptomic data with cellular resolution, e.g., 10x Xenium, can also provide accurate cell type information for each cell, and thus Concordia can be applied to such datasets. However, such spatial transcriptomic data are still too expensive for large-scale studies.

For each tissue section, we construct a **basic graph** by representing cells as nodes and connecting cells within a pre-defined Euclidean distance. We define the neighborhood of a cell as its 2-hop neighborhood in the basic graph, and define the neighborhood cell type composition (CTC) of a cell by the cell type proportions among all cells in its 2-hop neighborhood. To better capture complex patterns in tissues, we augment the basic graph through a two-step procedure. First, for each cell, we add edges to up to four cells within an enlarged search radius (3x the distance threshold of the basic graph) that have the most similar neighborhood CTC. We refer to this graph as the **local graph**. Second, we connect any two cells if there exists at least one path between them, they share similar neighborhood CTC, and most cells along the shortest path between them have similar neighborhood CTC. We refer to this graph as **extended graph** (Fig. 1A).

**Figure 1:**
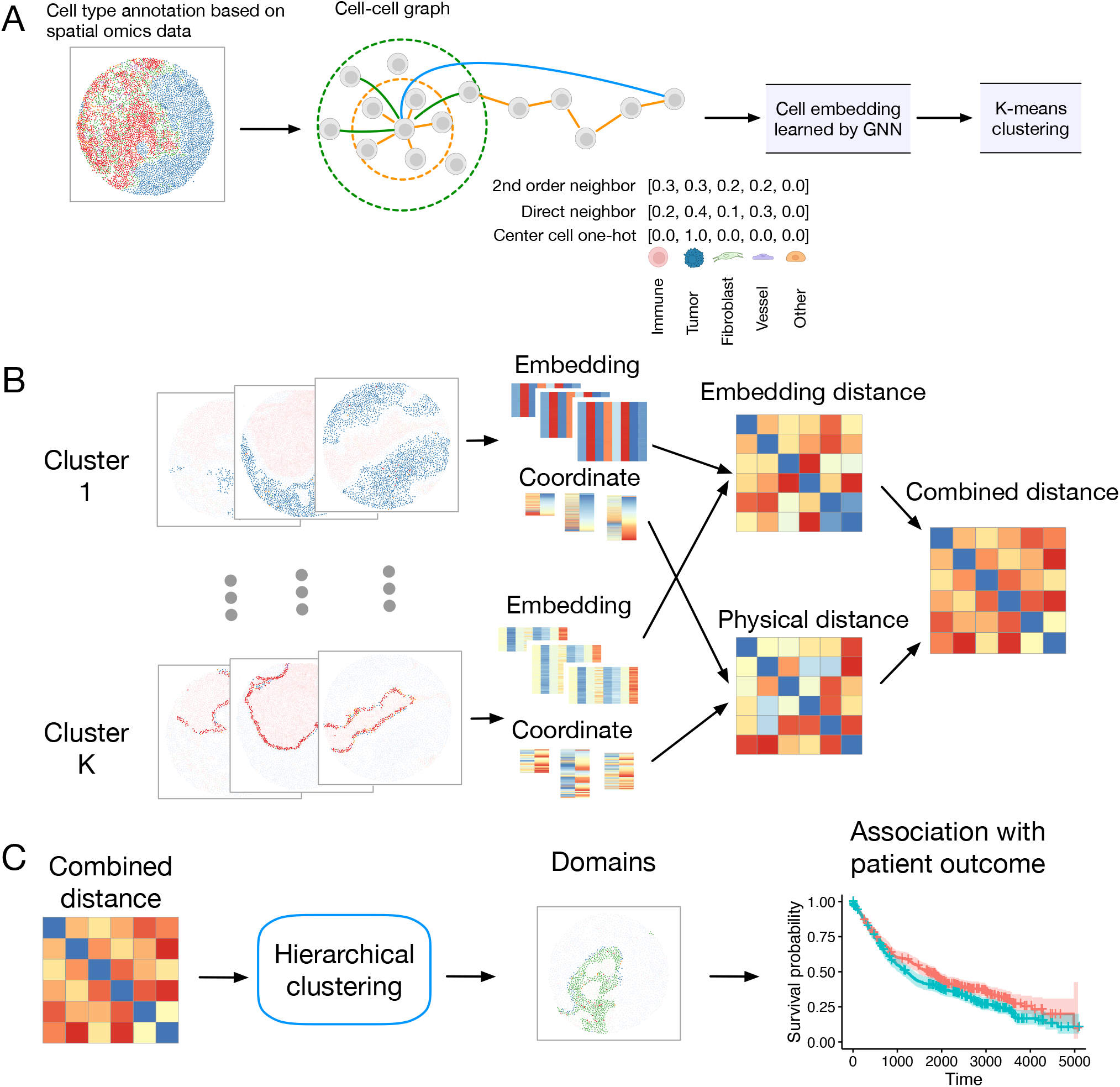
An overview of Concordia. (A) A cell-cell graph is constructed by connecting cells within certain distance threshold, which is subsequently expanded into an augmented graph where two cells are connected if they share similar cellular neighborhood and most of the cells along the path connecting them also share similar cellular neighborhood. The feature vector of a cell is generated by concatenating the center cell’s one-hot encoded cell type identity with its 1st and 2nd order neighborhood compositions. A GNN is trained with an unsupervised objective, and per-cell embeddings are extracted from the trained model. Cells are then clustered based on GNN-inferred embeddings.(B) Calculation of the distance between any two clusters obtained from step (A). The distance is a weighted combination of the distance in the embedding space and the distance by spatial coordinates. (C) The clusters obtained from step (A) are grouped into domains based on the distances obtained from step (B). These domains are subsequently used for downstream analyses, including associations with clinical outcomes. The cell type labels in panel (A) were created with BioRender.com.

The input feature vector for each cell is a concatenation of three components: (1) a one-hot encoding of the cell’s own cell type, (2) the CTC of its direct neighbors, and (3) the CTC of its second-order neighbors (Fig. 1A). This tri-level characterization provides a comprehensive description of the local cellular niche. These input feature vectors pass through the GNN to aggregate information across neighboring cells to generate new cell embeddings. After training the GNN, we extract the cell embedding and perform domain discovery in two steps (Fig. 1(B, C)). First, we apply *K*-means to cluster cells based on these cell embeddings and intentionally generate a larger number of clusters. Second, we quantify the distance between any two clusters by integrating the distance in the embedding space and the distance in the physical space, and then use this distance metric to perform hierarchical clustering and merge clusters into domains. We then leverage the inferred spatial domains in downstream analyses, for example, to assess associations with patient clinical outcomes (Fig. 1C).

For the GNN architecture, we use a graph attention network GATv2^20^ to perform graph convolution, allowing the model to dynamically weight neighbor importance. The network consists of two graph convolutional layers with a skip connection bridging the input to the output of the second layer (Fig. 2A). To enable unsupervised learning of cell clusters, the final embeddings are passed through a linear layer to produce a soft assignment matrix **S**.

**Figure 2:**
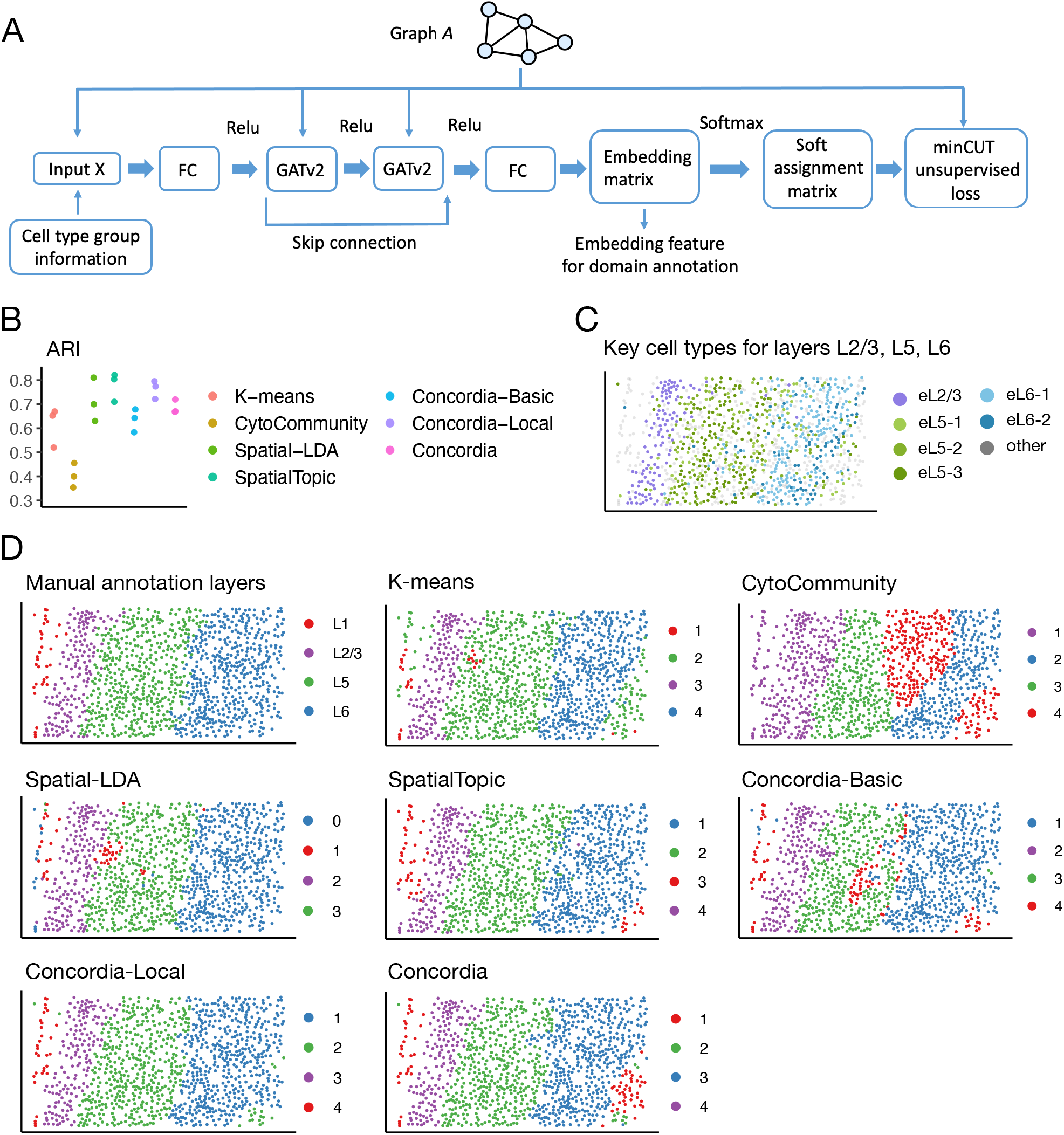
(A) Model architecture and domain annotation performance on mouse brain data. The graph is involved in computing the input cell feature vector, the two graph attention layers, and the unsupervised loss. The embedding feature for K-means clustering step is extracted after the second fully connected layer and before the softmax activation. FC: fully connected layer. GATv2: graph attention networks v2. (B) Domain annotation accuracy on the mouse brain dataset measured by adjusted Rand index (ARI) against manual domain labels (n=3 samples; each point is one sample; higher is better). A representative sample is shown with distributions of key excitatory neuron cell types for layers L2/3, L5 and L6 highlighted, manual layer annotation labels, and domain annotation from each method.

Each row of **S** corresponds to a cell and each column of it corresponds to a cluster. **S**_*ij*_ scores how likely the *i*-th cell belongs to the *j*-th cluster, and Σ_*j*_ **S**_*ij*_ = 1 for any *i*. We optimize the model using the unsupervised loss function minCUT,^21^ which encourages the partitioning of the graph into densely connected disjoint components. For datasets with multiple samples, a single model is trained jointly across all images with shared parameters, enabling learning of domain patterns that are comparable across samples within the same dataset. Although the soft assignment matrix **S** provides a direct way to cluster cells through a row-wise argmax operation (i.e., find the cluster with maximum weight), this approach does not capture the relative confidence of assignments. Instead, we use the learned cell embedding (Fig. 2A) and take the aforementioned two-step procedure to obtain spatial domains. To scale to cohort-sized spatial proteomics studies with large graphs, we implement minCUT using sparse adjacency matrices in PyTorch and use sparse message passing throughout training.

Among existing methods for spatial domain detection, CytoCommunity^19^ is the most closely related to Concordia. CytoCommunity is a GNN based method that performs unsupervised domain annotation on a *k*-nearest neighbor (kNN) cell graph. It has the same minCUT objective function as Concordia and optimizes it using the soft assignment matrix **S**. There are two major differences between the two methods. First, Concordia uses an extended cell-cell graph to allow connections of distant cells. Second, while both methods employ a minCUT loss to train the GNN, Concordia uses the resulting cell embeddings for downstream domain delineation, whereas CytoCommunity directly uses the learned soft assignment matrix **S** to assign a cell to the cluster with the highest weight (i.e., row-wise argmax operation on **S**).

### Concordia correctly identifies domains in well-structured tissues

Many earlier studies on spatial domain detection have focused on well-structured tissues, such as the brain, because these tissues exhibit clear anatomical organization. For example, the cerebral cortex is organized into distinct layers (L1–L6), providing a natural benchmark for evaluating spatial domain inference methods. While our ultimate goal is to characterize more complex and heterogeneous spatial domains such as those observed in tumors, we first demonstrate Concordia on cortical brain tissue, where well-defined cortex layers provide a principled and interpretable benchmark.

We evaluated Concordia on a publicly available mouse brain spatial transcriptomics dataset generated using the STARmap platform, consisting of three medial prefrontal cortex (mPFC) tissue sections from different mice.^22^ All cells were annotated to one of the 15 cell types, including subtypes of excitatory neurons, inhibitory neurons, and glial cells. As for the underlying true domain, we used the manual annotations provided by Li et al.,^6^ in which cells were assigned to four laminar domain labels (L1, L2/3, L5, and L6). Layer L4 was not included because mouse mPFC lacks a distinct L4 layer.^22, 23^

We compared Concordia with four existing methods. The first was *K-means* that clusters cells across images using the cell type composition of the 2-hop neighborhood of each cell as input features. The second was *CytoCommunity* .^19^ In addition, we also included two topic model-based methods, Spatial LDA^16^ and Spatial Topic.^17^ For Concordia, besides the extended graph, we evaluated the variants of Concordia that used the basic and local graphs, referred to as Concordia-Basic and Concordia-Local, respectively.

To ensure a fair comparison, all methods were configured to resolve four spatial domains. For the K-means method, we set the number of clusters to *K* = 4. For Concordia and its alternative options with simplified graphs (*basic* and *local*), we used an embedding dimension of *H* = 40, first performed clustering with *K* = 16 clusters, and then merged the 16 clusters into 4 domains. For CytoCommunity,^19^ we set the number of domains to 4 and otherwise followed the default parameter settings recommended by the authors. For Spatial-LDA^16^ and SpatialTopic,^17^ the topics are equivalent to domains for domain annotation, and are thus set to 4. We evaluated three values of radius parameter and presented the best performing results.

To quantify the accuracy of each method, we calculated the Adjusted Rand Index (ARI) between the predicted spatial domains and the ground-truth layer annotations for each tissue section. ARI measures the similarity between two clusterings with higher values indicating greater agreement. Fig. 2 shows the ARI comparison across all three tissue sections (B), the distributions of representative excitatory neuron subtypes across layers L2/3, L5, and L6 (C), as well as the manual layer labels alongside clustering results from the 6 methods for tissue section BZ14 (D).

Concordia and its two variants achieved comparable or higher ARI than *K-means* and CytoCommunity, while CytoCommunity showed the lowest agreement with the ground truth. Concordia-Local, Spatial-LDA and SpatialTopic performed the best with comparable ARI values. All three Concordia methods correctly identified L1 layer. In contrast, CytoCommunity mixed L1 and L2/3, and K-means mixed L1 and L5. The fact that K-means mixed two non-adjacent domains shows a key limitation of K-means: it does not incorporate spatial location information. The relative strong performance of Spatial-LDA and SpatialTopic is likely due to their design, which encourages spatially close cells to be annotated into the same domain, an appropriate assumption for well-structured tissues such as cortical layers. Similarly, Concordia with the local graphic networks also performed well under this setting. Domain annotation results for the other two tissues showed similar trends. Corresponding figures for these tissue sections are shown in Supplementary Figs. 1 and 2. Illustration of graph extensions are also included in these figures and in Supplementary Fig. 3.

### Concordia identifies meaningful domains in architecturally disorganized tissues

We applied six methods (*K-means, Spatial-LDA, SpatialTopic, Concordia-Basic, Concordia-Local*, and *Concordia*) to a lung cancer spatial proteomics dataset.^3^ This dataset comprises Imaging Mass Cytometry (IMC) images from tumor invasive-front cores. After quality control, 1,534 images from 860 patients were retained. We did not include CytoCommunity due to its relatively limited performance on well structured mouse brain data. The original study annotated 30 refined cell types. We grouped them into five coarser groups: immune, tumor, CAF (cancer-associated fibroblast), vessel, and other for our analysis. This consolidation is necessary because many refined cell types have very low proportions, making the graph sparse. For the three Concordia methods, we used an embedding dimension of *H* = 40 and first grouped all the cells into *K* = 40 clusters, which were subsequently merged into 10 domains. We chose 10 domains as a parsimonious resolution that captures the complex tumor characteristics, while maintaining within domain homogeneity. For Spatial-LDA^16^ and SpatialTopic,^17^ we also set the number of topics to 10.

In this real dataset, ground-truth spatial domains are unavailable. Therefore, agreement-based metrics such as the adjusted Rand index (ARI) cannot be used to evaluate domain annotations. Instead, we compared methods using two approaches. For methods that use per-cell numerical features (*K-means, Basic, Local*, and *Extended*), we evaluated the spatial dependence of these features and the extent to which they were explained by the inferred domains. For Spatial-LDA^16^ and SpatialTopic,^17^ which do not provide cell-specific embeddings like Concordia, we assessed their inferred domains by visual inspection and quantifying within-domain connectivity.

For the first approach, the feature representations for *K-means* are the cell type proportions in each cell’s 2-hop neighborhood, whereas for the three Concordia methods, the features are the embeddings learned by the corresponding GNN models. To evaluate spatial dependence, we measured whether spatially close cells tended to have similar features, quantified by Moran’s *I*. For each feature dimension, Moran’s *I* was computed separately within each image and then summarized by the median across images. Next, we assessed how well the inferred domains explain the variation of features. In other words, whether cells assigned to the same domain tended to have more similar features than cells assigned to different domains. We quantified this by an *R*^2^ value for each feature dimension, obtained from a linear model with the feature value as the response and domain membership as a categorical predictor.

The embedding features from the Concordia methods exhibited higher median Moran’s *I* than the neighborhood composition features used by *K-means*(Fig. 3A), consistent with the design of Concordia (i.e., the underlying GNN) that encourages similarity among connected nodes in the input graph. Both Concordia-Local and Concordia had higher *R*^2^ than K-means, whereas Concordia-Basic had slightly lower *R*^2^ (Fig. 3B). The *R*^2^ of K-means features were notably more variable across features than those of the Concordia methods.

**Figure 3:**
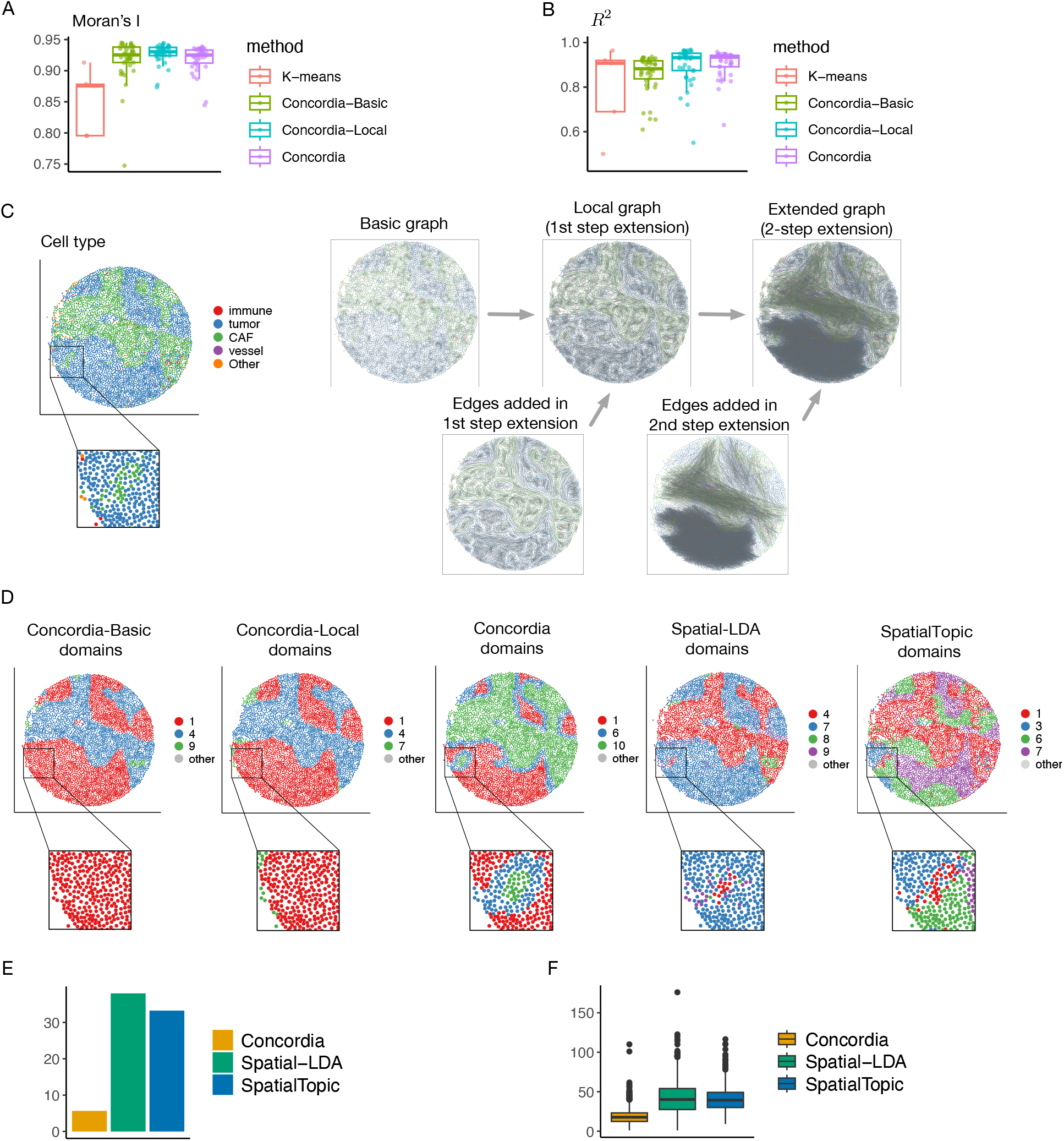
Comparisons of different methods in a lung cancer dataset.^3^ (A) Comparison of Moran’s I where higher values indicate nearby cells have more similar features. Here each point is the median of one embedding feature across all images. (B) Variance explained (*R*^2^) by identified domains for each feature. There is only one value below 0.5 (*R*^2^ = 0.2 for one feature of K-means) and it is clipped to 0.5 for better visualization. (C) Illustration of graph extension steps of Concordia on one image, including three types of graphs. (D) Illustration of domain annotations given by five methods both on the full image and the zoom-in region in Panel (C). Only domains that have at least 30 cells in the image are highlighted in each panel. Other domains are combined and labeled as “other”. (E)-(F) Number of within-domain connected components averaged across domains. (E) Barchart for the values in the current image. (F) Boxplots for the values from all images.

For the second approach, we examined representative domain annotations and quantified the spatial connectedness of the inferred domains. To illustrate the effect of graph augmentation, we examined the basic graph, the local graph, and the extended graph for a few representative lung cancer invasive front sections (Fig. 3C, Supplementary Fig. 4A). In the example shown in Fig. 3D, domain annotations by the default Concordia recover a distinct tumor-border region adjacent to CAF (i.e., the tumor cells at the tumor–CAF interface), which is not resolved when using Concordia-Basic, Concordia-Local, Spatial-LDA or SpatialTopic. Concordia also identifies a small group of CAFs surrounded by tumor cells (highlighted in the zoom-in regions of Fig. 3C,D), whereas this structure is missed by the Concordia-Basic or Concordia-Local. Although Spatial-LDA and SpatialTopic capture portions of this CAF cluster, neither method delineates the boundary as clearly as Concordia. A second example shows a similar pattern that Concordia, which uses the two-step augmented graph, recovers more refined domains than the alternative methods (Supplementary Fig. 4A,B).

To quantify the spatial connectivity of inferred domains, in each image, we computed the number of connected components within each domain and averaged this quantity across domains. Two cells are considered connected if they are neighbors in the basic graph. For a given domain, the connected components are counted based on the induced subgraph containing only cells within the same domain. Lower values indicate that cells assigned to the same domain form more spatially contiguous structures. We compared three methods, Concordia, Spatial-LDA and spatialTopic. In the example image shown in Fig. 3C, Concordia yielded a substantially lower average number of within-domain connected components than Spatial-LDA and spatial Topic (Fig. 3E). This is consistent with the visual observation from the domain annotation figures (Fig. 3D), that Concordia shows smoother and more connected shapes of annotated domains. This pattern holds across all images (Fig. 3F).

### Spatially defined domains identify cell subsets associated with clinical outcomes

We applied Concordia to two datasets: lung cancer and breast cancer. For both datasets, we set the number of domains to be 10. Because our model is trained jointly across all images, it yields spatial domains with consistent semantics, enabling cohort-level association analyses with clinical outcomes. Specifically, for each image, we calculated domain proportions as the fraction of cells assigned to each domain. For patients with multiple images, proportions were averaged across images. We assessed the marginal association of domain proportions with overall survival using a Cox proportional hazards model. We also additionally adjusted for clinical covariates (specified below for each cohort). A two-sided p-value less than 0.05 is considered significant. For interpreting domains, we examined cell type composition within each domain.

#### Lung cancer cohort

Domain 10, which was dominated by CAFs (Fig. 4A; Supplementary Fig. 2), showed a significant association with overall survival after Bonferroni correction (Fig. 4B). It remained significant after adjusting for age, sex, cancer type(LUSC and LUAD), grade (1, 2 and 3), and smoking status. Stratifying patients by the median Domain 10 proportion yielded significantly different Kaplan-Meier survival curves (log-rank *p* = 0.016; Fig. 4C). In contrast, the overall proportion of CAF cells was not significantly associated with survival (Fig. 4B). This suggests a functional divergence among CAFs depending on their spatial niche. When we stratified the CAF population by domain membership, only CAFs residing within Domain 10 showed a significant association with survival, whereas CAFs outside this domain did not (Figure 4D). Since the cell types in this dataset were defined by protein expression, this finding indicates that domain membership identifies a spatially defined CAF subset with distinct clinical relevance, which would be missed when examining protein expression only.

**Figure 4:**
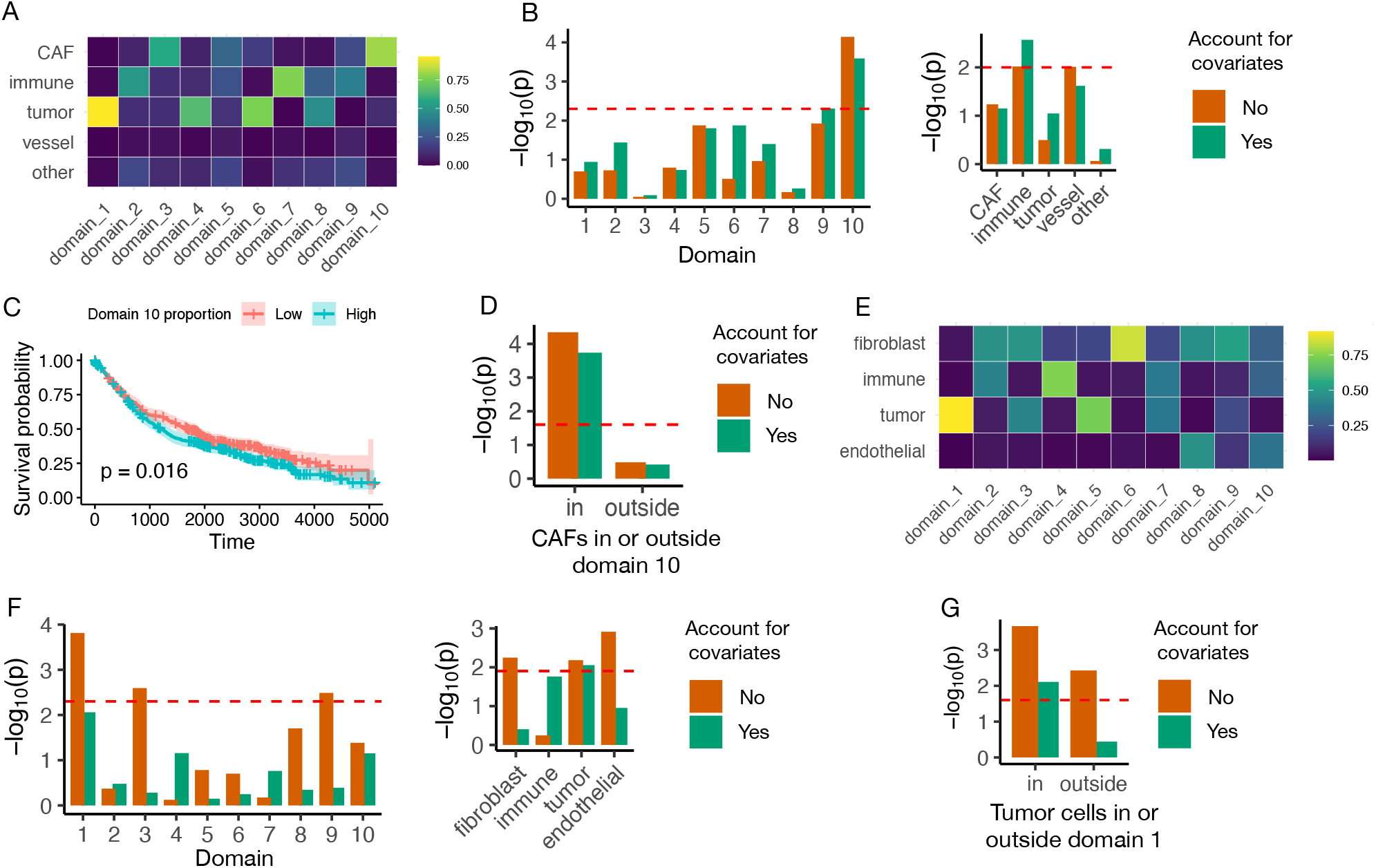
Spatial domain characterization in lung and breast cancer cohorts. (A-D) Lung cancer dataset analysis. (A) Median cell type proportions for each of 10 domains identified by the default Concordia using the “extended” graph. Here median is computed across images. (B) Associations between overall survival and Concordia domain proportions (left) or cell type proportions (right). Two types of Cox models were evaluated, either with or without covariates: age, sex, cancer type, grade, and smoking status. Dashed lines indicate Bonferroni-corrected significance thresholds. (C) Kaplan-Meier survival curves stratified by median proportion of domain 10 from panel (A). (D) Cox association p-values for CAFs within or outside domain 10 (Bonferroni-corrected *α* = 0.05*/*2). (E-G) Breast cancer dataset analysis. (E) Median celltype proportions within each of 10 domains identified by the default Concordia method. (F) Cox association p-values for each domain (left) or the proportion of each cell type (right) using marginal and covariate-adjusted models (covariates: age, grade, tumor stage, and PAM50 subtypes).(G) Cox association p-values for proportion of tumor cells within or outside domain 1. The dashed line indicate Bonferroni-corrected significance threshold (*α* = 0.05*/*2).

#### Breast cancer METABRIC cohort

We also analyzed a breast cancer IMC dataset derived from METABRIC,^4, 24^ in which pathologist-selected invasive cancer regions were imaged. After quality control, 488 images from 549 patients were retained. We identified a tumor-dominated domain (Domain 1; Fig. 4E and Supplementary Fig. 6) that was significantly associated with overall survival after Bonferroni correction (Fig. 4F); however, this association became not significant after adjusting for age, grade, tumor stage, and PAM50 subtypes. The overall tumor-cell proportion was also significantly associated with survival. Partitioning tumor cells into those inside versus outside Domain 1 showed that tumor cells within Domain 1 showed a stronger association with survival than those outside (Fig. 4G), suggesting the existence of a spatially defined tumor-cell subset of tumor cells with distinct prognostic relevance.

## Discussion

We present Concordia, a GNN-based framework for unsupervised spatial domain annotation that jointly models cell-type labels and spatial coordinates. The model leverages an augmented cell-cell graph designed to capture complex tissue structures such as tumor invasive fronts that would be missed by methods relying on distance-threshold graphs. Rather than partitioning the graph directly, Concordia derives domains from learned cell embeddings by a two-step approach: first over-clustering the cells and then merging clusters into domains. This approach improves robustness of domains by decoupling representation learning from the choice of the number of domains. The resulting domains exhibit smoother shapes and better within-domain connectivity than other methods such Spatial-LDA and SpatialTopic. Importantly, Concordia trains a single model across all images, yielding consistently defined domains across images that enable cohort-level analyses. Applied to two real datasets, Concordia identified domains with distinct clinical relevance. The choice between Concordia and its local graph variant, Concordia-Local, depends on the structure characteristics of the tissue. Concordia-Local is well suited for regular, well-organized structures (e.g., cortical layers in brain), whereas Concordia is preferred for tissues involving complex shapes with elongated or branching geometries (e.g., tumor invasive front).

Spatial proteomics data can be analyzed at the level of protein marker abundance or inferred cell types. Directly analyzing protein abundance can help identify subtle cell states. However, for population-level analysis, using inferred cell types is more robust to batch effect, more interpretable, and thus better suited for our purpose for identifying consistently defined spatial domains. An additional benefit is that cell type based analysis transfer more naturally across datasets. This is particularly valuable when marker panels differ across cohorts, as cell type labels provide a shared taxonomy that numerical molecular measurements cannot.

Several questions remain open. First, the choice of granularity of cell types may influence both graph construction and biological interpretation. Coarser cell types (e.g., immune, tumor, CAF, vessel) increase the number of candidate edges during graph augmentation by making neighborhood similarity more likely to exceed the threshold, improving the ability to capture long-range or fragmented structures. They also allow identification of subsets defined jointly by coarse cell type calls and spatial location information, simplify interpretation and facilitate cross-dataset comparisons. However, overly coarse cell types may obscure biologically meaningful heterogeneity. For example, lumping CD4+ and CD8+ T cells into a single immune category could mask domain differences in their spatial organization. Developing principled strategies for cell type grouping, potentially data-adaptive and context-dependent, is an important direction for future work.

Second, the choice of the number of domains also affects the resolution of the annotation. This mirrors the classical challenge of choosing the number of clusters in an unsupervised learning, where challenges and solutions also apply here. The number of domains can be selected based on prior knowledge of tissue organization (e.g., four cortical layers in brain) or guided empirically by the similarity of spatial clusters within domains, e.g., evaluating the within-domain distances as a function of the number of clusters until it is flatten (i.e., the so-called elbow method in K-means clustering).

Third, while our sparse implementation of the minCUT loss improves scalability of model training, the computational cost of graph augmentation remains a consideration. In particular, the second graph augmentation step is computationally demanding because it requires evaluating the quality of shortest-paths between many candidate cell pairs. Improving the efficiency of this step, for example, by selectively choosing the candidate cell pairs to consider before evaluating the shortest-paths, could further extend applicability to even larger images and cohorts.

In terms of computational resources, GNN model training was run on a NVIDIA L40S GPU with 48GB memory and all other steps were run on Intel Xeon Gold 6254 CPUs (3.10GHz) on a computing server. On the lung cancer dataset, for a typical image with the median number of cells (around 3,080), generating the extended graph took about 1.5 minutes. When parallelized across 30 cores, generating extended graphs for the entire dataset of 1,534 images could be completed within 2 hours. GNN model training took around 3 hours. The final processing steps after GNN model training took 20 minutes in total: this included running k-means clustering on the learned embeddings, followed by the between-cluster distance matrices computation per image, and finally obtaining the domains. Among them, the step of distance matrices computation was completed in 5 minutes when parallelized across 30 cores.

Our method focuses on spatial proteomics data, where molecular measurements are typically more restricted than in spatial transcriptomics. It can also be applied to single-cell level spatial transcriptomics data with cell type annotation. In spatial transcriptomics, additional sources of information, such as gene programs and inferred cell–cell communication, may provide further resolution for domain annotation.^25, 26^ Incorporating these signals is beyond the scope of the present study. Nevertheless, the graph augmentation strategy introduced here could potentially be useful to spatial transcriptomics data as well, though additional work will be needed to define neighborhood similarity. Unlike in proteomics, gene expression captures not only cell types but also dynamic cell states, making the definition of neighborhood similarity that is both robust and biologically meaningful a non-trivial challenge.

## Methods

### Computational models

Our method operates on spatial single-cell datasets with (i) per-cell coordinates and (ii) discrete cell-type labels. When datasets contain a large number of fine-grained cell types, we optionally merge them into a smaller set of coarser cell types (e.g., grouping B cells and T cells as “Immune”) to improve robustness and reduce feature sparsity. For each tissue section (image), we construct an undirected spatial graph (**basic graph**) *G*_0_ = (*V, E*_0_) in which nodes represent cells and edges connect pairs of cells whose Euclidean distance is below a threshold *r*. To choose the value of *r*, we randomly sample a subset of images and selected the distance cutoff *r* such that the median (across sampled images) of the resulting graph’s average node degree was approximately 6. Using the basic graph *G*_0_, we compute for each cell *i* a 2-hop neighborhood composition vector *c*_*i*_, defined as the normalized counts of cell types among nodes within two hops of cell *i* in the graph. Here, normalization refers to converting counts into proportions such that the entries of *c*_*i*_ sum to 1. The vector *c*_*i*_ is called the neighborhood cell type composition (neighborhood CTC) for cell *i*.

#### Graph augmentation

Starting from *G*_0_, we form an augmented graph via two extension steps. Step 1 (local gap-bridging). For each cell *i*, we search candidate neighbors within an enlarged spatial neighborhood (within 3\**r* radius) and add edges to the top 4 most similar candidates according to Euclidean distance ∥*c*_*i*_ − *c*_*j*_ ∥_2_. This step introduces additional local connections to improve connectivity across narrow gaps. The resulting graph is **local graph**. Step 2 (long-range connection). We further add edges between spatially distant cells *i* and *j* when they are connected in the Step-1 graph (local graph) by a shortest path whose intermediate nodes predominantly share similar neighborhood composition (measured by ∥ · ∥ _2_ on *c*) with both cells *i* and *j*. To control graph density and computational cost, we subsample eligible long-range pairs to match a target average node degree (20 by default) in each graph, with sampling biased toward pairs with longer shortest-path distances to preferentially add long-range edges. The resulting graph is **extended graph**. More details on the two steps of graph extensions, including the parameters for settings the requirements on the neighborhood CTC, are in Details of methods section in Supplementary Materials.

#### Input and loss function for graph neural network

The GNN operates on a dual-input architecture consisting of the augmented cell-cell adjacency graph and a feature matrix (Fig.2A). For each cell *i*, we define a composite feature vector *x*_*i*_ ∈ ℝ^3*C*^, where *C* denotes the total number of unique cell types. This vector is constructed through the concatenation of three distinct descriptors, capturing the hierarchical neighborhood structure: (1) A one-hot encoded vector representing the specific cell type of the center cell; (2) A composition vector representing the normalized distribution of cell types among the cell’s immediate (direct) neighbors; and (3) A composition vector derived from the cell’s exclusive second-order neighbors (nodes reachable in exactly two hops, excluding direct neighbors).

We employ a GNN to perform message passing, iteratively aggregating contextual features from the neighborhood into the center cell representation (Fig. 2A). Specifically, we use GATv2 graph convolution,^20^ which implements a dynamic attention mechanism that allows the model to weigh the importance of specific neighboring nodes. Our architecture consists of two graph convolutional layers. To mitigate the risk of over-smoothing and facilitate optimization, we incorporate a residual skip connection that bypasses the convolutional layers, directly linking the input to the first GATv2 layer to the output of the second GATv2 layer (prior to the ReLU nonlinearity). The resulting node representations are passed through a linear projection and a softmax to obtain a soft assignment matrix **S** ∈ ℝ^*N ×H*^, where *N* is the number of cells and *H* is the number of components. Each row sums to 1 and can be viewed as encoding the soft membership of the cell across the *H* components. Model parameters are optimized using the minCUT objective,^21^ which is grounded in spectral graph theory, aiming to partition the graph into disjoint, densely connected components by minimizing the cut of the edges between them while ensuring balance in the cluster sizes. For a single image, given the adjacency matrix **A** and the soft assignment matrix **S**, the formula for the minCUT loss function is:

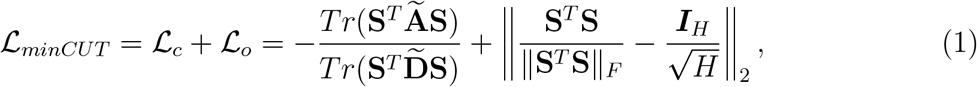

where **Ã** is the symmetrically normalized adjacency matrix, i.e., 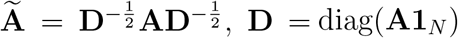, and 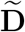 is the diagonal matrix for **Ã** . In addition, *H* is the number of columns of the soft assignment matrix **S**, and ∥ · ∥_*F*_ is Frobenius norm. Minimizing the first part of the loss function, ℒ_*c*_, encourages nodes strongly connected in the graph to have similar representations, and minimizing the second part of the loss function, the orthogonality loss term ℒ_*o*_, encourages the cluster assignments to be orthogonal and the clusters to be of similar size.

To ensure a robust and consistent representation of spatial domains across a cohort, our model is trained through joint optimization across all images in the dataset. The training objective is defined as the sum of per-image minCUT losses, i.e., 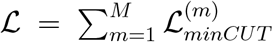 where *M* is the number of images and 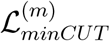 denotes the loss computed on image *m*. By employing global parameter sharing, the GNN architecture maintains a unified set of weights across all spatial graphs. This enables the model to learn a generalized embedding space that is invariant to sample-specific noise, facilitating the identification of consistent biological patterns across heterogeneous tissue sections.

#### Obtain domain annotation

Because the minCUT objective is defined in terms of the soft assignment matrix **S**, a straightforward strategy is to set the number of columns of **S** to the desired number of domains *H* and convert **S** to hard label by assigning each cell to argmax_*h*_**S**_*ih*_. However, this hardening step discards assignment confidence and treats highly confident and weakly assigned cells equivalently. To mitigate the inconsistencies associated with hard assignments derived directly from **S**, we use the pre-softmax activations as per-cell embeddings (Fig. 2A). We adopt a two-stage procedure that first identifies a larger number of spatial clusters and then merges them into a smaller number of spatial domains using both embedding and physical (coordindate) information.

Step 1 (overclustering in embedding space). We pool cells from all images and apply *K*-means to the extracted embeddings to obtain *K* clusters; by default we use *K* = 40 to allow sufficient granularity. Because the GNN parameters are shared across all images, these 40 clusters are semantically consistent across the entire dataset.

Step 2 (merging clusters into domains). To merge the *K* embedding-derived clusters into domains, we compute two inter-cluster distance measures for each cluster pair (*a, b*), aggregating evidence across images that contain both clusters with enough number of cells (qualified images).

Embedding-space distance. Within each qualified image *m* (image *m* is considered qualified for a cluster pair (*a, b*), if both clusters occupy a sufficient number of cells within it), we compute the Euclidean distance between the embedding centroids of clusters *a* and *b*, 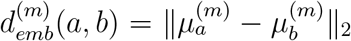. The embedding distance at the data set level is defined as the median across qualified images:

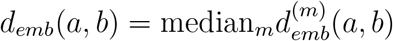

Physical (spatial) distance. Within each qualified image *m*, we quantify spatial separation using directed nearest-neighbor distances between clusters. For each cell *i* in cluster *a*, let *δ*(*i, b*) denote the Euclidean distance from *i* to its nearest cell in cluster *b*. We compute the 90th percentile of these distances over cells in *a*, denoted 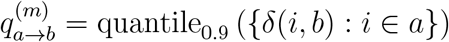 and analogously 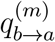 for cells in *b*. The image-level physical distance is then

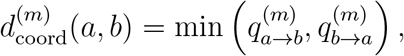

and the dataset-level physical distance is the median across qualified images:

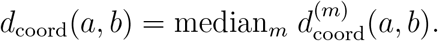

For each cluster pair, we aggregate the embedding distances and physical distances across images separately, and obtain two distance matrices, one capturing embedding-space proximity and the other capturing physical spatial proximity, where each entry corresponds to a specific pair of clusters. To place the two metrics on a comparable scale, we rescale the embedding-distance matrix by a scalar chosen so that its median over all pair of clusters matches the median of the physical-distance matrix. We then fuse the two matrices into one by taking their arithmetic mean and apply hierarchical clustering to obtain the final domain assignments.

More details on merging *K*-means clusters into domains, including the parameter for deciding whether an image is qualified for two given clusters and how to assign between-cluster distance when two clusters not occurring together in enough number of images, are in Details of methods section in Supplementary Materials.

#### Scalability

Existing implementations of the minCUT objective are typically expressed in terms of a dense adjacency matrix, which can be memory- and compute-prohibitive for large spatial graphs. We therefore implement minCUT to accept sparse adjacency matrices in the COO format, enabling loss computation to scale with the number of edges *E* rather than the number of node pairs *N*^2^. Because the same graph is used for both the loss and GNN message passing, this sparse representation also allows the graph convolution layers to operate efficiently on large images and cohort-scale datasets. In our implementation, the dominant dense operations that scale quadratically in *N* are replaced by sparse matrix operations whose cost is proportional to *E* (with additional terms depending on the number of components *H* and feature dimension *F*).

#### Differences from CytoCommunity

Our approach is most closely related to CytoCommunity,^19^ which also performs unsupervised domain annotation using a cell graph, a GNN, and the minCUT objective. The key differences are: (i) graph construction, and (ii) domain assignment strategy.

1. **Graph construction**. CytoCommunity uses a *k*-nearest-neighbor (kNN) graph. We begin with a distance-threshold (radius) graph and augment it by adding edges guided by 2-hop neighborhood composition similarity and by qualified long-range connections. This allows our model to capture higher-order architectural motifs and resolve complex microenvironmental boundaries that proximity-based graphs may overlook.
2. **Domain assignment**. CytoCommunity converts the minCUT soft assignment matrix to hard labels and stabilizes results via an ensemble (majority voting across multiple runs). In contrast, we extract pre-softmax embeddings from a single trained model, overcluster cells by *K*-means, and merge clusters into domains using a combined embedding–spatial distance. This avoids the information loss associated with direct argmax discretization of the soft assignment matrix.

### Evaluation metrics

#### Moran’s *I*

Moran’s *I* is computed with Python package `Scanpy`. For each feature dimension, we compute Moran’s *I* within each image and summarize results by the median Moran’s *I* across all images. Here the Moran’s *I* spatial weights are derived from physical proximity (coordinate *k*NN computed with function `scanpy.pp.neighbors`, with the default `n_neighbors=15` specified by the function), independent of the augmented graphs used during GNN training.

#### R^2^

The *R*^2^ is computed on a set of 100 randomly chosen images from the lung cancer dataset. The sample unit in the linear regression is each cell in the 100 images. For each embedding (or feature) dimension, the *R*^2^ value is obtained from fitting a linear regression *y* ∼ *domain*, where *y* is the value on the given dimension, *domain* is the categorical domain annotation given by the method. The computation of *R*^2^ was done in `R` version 4.4.0.

#### Average number of connected components

To compute the number of connected components, for each image, for each method, based on the domain annotation, we only consider the domains that each has at least 30 cells in the given image. Within each domain, making use of the basic graph, the number of connected components is computed based on the subgraph induced by cells in this domain. The Python function used for computing the number of connected components is `networkx.number_connected_components` from the package `networkx`. For each image, the final number is the average of the numbers of connected components from all qualified domains.

### Comparing to other methods

#### CytoCommunity (v1.1.0)

CytoCommunity (unsupervised mode) was run and evaluated only on the mouse mPFC dataset. We followed the approach suggested in the original paper and set the number of *K* in KNN as 32, which is around the squared root of the number of cells in a sample (around 1,000).

#### Spatial-LDA (v0.1.3)

On the mouse mPFC dataset, we ran Spatial-LDA using radius parameter values 490, 690, and 1600, corresponding to each cell on average having around 10, 20, and 100 neighbors within the radius. On the lung cancer dataset, we ran Spatial-LDA using radius parameter values 27 and 62, corresponding to each cell on average having around 20 and 100 neighbors within the radius. For each cell, from the resulting weights of each cell for all topics, we picked the topic with the highest weight as the assigned domain of the cell. On the mouse mPFC dataset, the best performances in terms of ARI are from radius 690 and the corresponding results are shown in Fig. 2B, Supplementary Fig. 1A and Supplementary Fig. 2A. The ARI values of all radius parameter settings are included in Supplementary Table 1. On the lung cancer dataset, the results from using radius 27 are shown in Fig. 3 while the run with radius 62 failed in the optimization process.

#### SpatialTopic (v1.2.0)

On both mouse mPFC and lung cancer datasets, we ran SpatialTopic using the same radius settings as used for running SpatialTopic. The corresponding sigma parameter is set to be around square root of the radius, following the suggestion in the original paper. For each cell, we directly used the topic output by SpatialTopic as the assigned domain of the cell. On the mouse mPFC dataset, the best performances in terms of ARI are from radius 690 and the corresponding results are shown in Fig. 2B, Supplementary Fig. 1A and Supplementary Fig. 2A. The ARI values of all radius parameter settings are included in Supplementary Table 1. On lung cancer dataset, we show the results from the visually more appealing setting radius 62 in Fig. 3 and Supplementary Fig. 4. The results from using radius 27 show similar patterns and are included in Supplementary Fig. 7.

### Datasets

#### Lung cancer dataset

The lung cancer dataset was downloaded following the resource provided in original work.^3^ We restricted the analysis to patients satisfying the following critera: (1) having a documented relapse status within 15 years, (2) having a diagnosis of either adenocarcinoma or squamous cell carcinoma, and (3) not having received neoadjuvant therapy. For the spatial proteomics data, two ROIs were excluded based on anomalous “Area mm Core” and “Area px Core” values that each is likely a mislabeling that result from combining two different ROIs. Further, we excluded ROIs with number of cells less than 1,000. After filtering steps, 1,534 ROIs from 860 patients were retained for model training.

The dataset has 30 cell types organized into six broad cell type categories: fibroblast, immune, T cell, tumor, vessel, and other that includes all cells not belonging to the aforementioned categories. We combine immune and T cell and consider this as a broad “immune” category and treat the rest the same. Specifically, “CAF” includes collagen_CAF, dCAF, hypoxic_CAF, hypoxic_tpCAF, iCAF, IDO_CAF, mCAF, PDPN_CAF, SMA_CAF, tpCAF, and vCAF. “Immune” includes Bcell, CD4, CD4_Treg, CD8, IDO_CD4, IDO_CD8, ki67_CD4, ki67_CD8, Myeloid, Neutrophil, PD1_CD4, TCF1/7_CD4, TCF1/7_CD8. “Tumor” includes hypoxic and normal. “Vessel” includes blood, HEV, and Lymphatic.

#### Breast cancer dataset

For the METABRIC breast cancer dataset, the image data and one clinical data table were downloaded from the resource provided in the original study.^4^ Another patient clinical data table with longer follow-up time was downloaded from cBioPortal. Most tissue spots were 0.6 mm in diameter, but one TMA block contained spots that were 1 mm in diameter. The single cell data object includes both data for both tumor and normal images. We only kept the tumor images that (1) have at least 1,000 cells and (2) are from patients with known ER status in the clinical data table from the original study, excluding 2 images from patients with unknown ER status. After quality control, 488 images from 549 patients were kept for model training.

There are 32 cell types in the dataset, which we grouped into four broad categories: immune, tumor, fibroblast and endothelial. “Immune” includes cell types T_Reg, T_Ex, CD4^+^ T cells & APCs, CD4^+^ T cells, CD8^+^ T cells, B cells, CD38^+^ lymphocytes, granulocytes, macrophages, macrophages & granulocytes, Ki67^+^, and CD57^+^. “Tumor” includes cell types MHC I & II^*hi*^, MHC I^*hi*^CD57^+^, MHC^*hi*^CD15^+^, HER2^+^, CK8-18^+^ ER^*hi*^, CK^*lo*^ER^*med*^, Basal, ER^*hi*^CXCL12^+^, CK8-18^*hi*^CXCL12^*hi*^, CD15^+^, Ep CD57^+^, CK^+^ CXCL12^+^, Ep Ki67^+^, CK^*med*^ER^*lo*^, CK^*lo*^ER^*lo*^, and CK8-18^*hi*^ER^*lo*^. “Fibroblast” includes cell types Fibroblasts, Fibroblasts FSP1^+^, Myofibroblasts, and Myofibroblasts PDPN^+^. The endothelial cell type itself is treated as its own category “endothelial”.

For survival analysis, we used overall survival information from the clinical data table obtained from cBioPortal. Among the clinical covariates included in the model, tumor grade and PAM50 subtypes are from the original study’s clinical table, while age and tumor stage are from the cBioPortal clinical table.

### Mouse mPFC dataset

For the mouse medial prefrontal cortex data by STARmap^22^,^6^ We use the version from a later study^6^ where the manually annotated layer labels were obtained based on spatial gene expression patterns and the Allen Mouse Brain Atlas. There are 15 cell types, Smc, Endo, Astro, eL6-2, Oligo, eL5-3, Reln, VIP, eL2/3, NPY, eL5-2, Lhx6, L5-1, SST, and eL6-1. Each of them is directly treated as a cell type in the model.

## Code availability

The version of mincut pool operator that takes sparse adjacency matrix as input for computing unsupervised loss is available on GitHub (https://github.com/Sun-lab/sparse_mincut_pool). The code files for running our methods are available on GitHub with a tutorial (https://github.com/Sun-lab/Concordia). Our data analysis pipeline is available at (https://github.com/Sun-lab/Concordia_pipeline).

## References

[1] Giesen C, Wang HA, Schapiro D, Zivanovic N, Jacobs A, Hattendorf B, et al. Highly multiplexed imaging of tumor tissues with subcellular resolution by mass cytometry. Nature methods. 2014;11(4):417–422.

[2] Goltsev Y, Samusik N, Kennedy-Darling J, Bhate S, Hale M, Vazquez G, et al. Deep profiling of mouse splenic architecture with CODEX multiplexed imaging. Cell. 2018;174(4):968–981.

[3] Cords L, Engler S, Haberecker M, Rüschoff JH, Moch H, de Souza N, et al. Cancer-associated fibroblast phenotypes are associated with patient outcome in non-small cell lung cancer. Cancer Cell. 2024;.

[4] Danenberg E, Bardwell H, Zanotelli VR, Provenzano E, Chin SF, Rueda OM, et al. Breast tumor microenvironment structures are associated with genomic features and clinical outcome. Nature genetics. 2022;54(5):660–669.

[5] Wu Z, Trevino AE, Wu E, Swanson K, Kim HJ, D’Angio HB, et al. Graph deep learning for the characterization of tumour microenvironments from spatial protein profiles in tissue specimens. Nature Biomedical Engineering. 2022;6(12):1435–1448.

[6] Li Z, Zhou X. BASS: multi-scale and multi-sample analysis enables accurate cell type clustering and spatial domain detection in spatial transcriptomic studies. Genome biology. 2022;23(1):168.

[7] Hu J, Li X, Coleman K, Schroeder A, Ma N, Irwin DJ, et al. SpaGCN: Integrating gene expression, spatial location and histology to identify spatial domains and spatially variable genes by graph convolutional network. Nature methods. 2021;18(11):1342–1351.

[8] Singhal V, Chou N, Lee J, Yue Y, Liu J, Chock WK, et al. BANKSY unifies cell typing and tissue domain segmentation for scalable spatial omics data analysis. Nature genetics. 2024;56(3):431–441.

[9] Long Y, Ang KS, Sethi R, Liao S, Heng Y, van Olst L, et al. Deciphering spatial domains from spatial multi-omics with SpatialGlue. Nature Methods. 2024;21(9):1658–1667.

[10] Wang J, Huo Y, Zhao R, Pan Y, Wang H, Li X. SpaMOAL: A spatial multi-omics graph contrastive learning method for spatial domains identification. bioRxiv. 2026; p. 2026–02.

[11] Kim J, Rustam S, Mosquera JM, Randell SH, Shaykhiev R, Rendeiro AF, et al. Unsupervised discovery of tissue architecture in multiplexed imaging. Nature methods. 2022;19(12):1653–1661.

[12] Wu Z, Kondo A, McGrady M, Baker EA, Chidester B, Wu E, et al. Discovery and generalization of tissue structures from spatial omics data. Cell Reports Methods. 2024;4(8).

[13] Qian J, Shao X, Bao H, Fang Y, Guo W, Li C, et al. Identification and characterization of cell niches in tissue from spatial omics data at single-cell resolution. Nature Communications. 2025;16(1):1693.

[14] Lundberg E, Borner GH. Spatial proteomics: a powerful discovery tool for cell biology. Nature Reviews Molecular Cell Biology. 2019;20(5):285–302.

[15] Hao Y, Stuart T, Kowalski MH, Choudhary S, Hoffman P, Hartman A, et al. Dictionary learning for integrative, multimodal and scalable single-cell analysis. Nature biotechnology. 2024;42(2):293–304.

[16] Chen Z, Soifer I, Hilton H, Keren L, Jojic V. Modeling multiplexed images with spatial-LDA reveals novel tissue microenvironments. Journal of Computational Biology. 2020;27(8):1204–1218.

[17] Peng X, Smithy JW, Yosofvand M, Kostrzewa CE, Bleile M, Ehrich FD, et al. Decoding Spatial Tissue Architecture: A Scalable Bayesian Topic Model for Multiplexed Imaging Analysis. bioRxiv. 2024;.

[18] Kipf T. Semi-supervised classification with graph convolutional networks. arXiv preprint arXiv:160902907. 2016;.

[19] Hu Y, Rong J, Xu Y, Xie R, Peng J, Gao L, et al. Unsupervised and supervised discovery of tissue cellular neighborhoods from cell phenotypes. Nature Methods. 2024;21(2):267–278.

[20] Brody S, Alon U, Yahav E. How attentive are graph attention networks? arXiv preprint arXiv:210514491. 2021;.

[21] Bianchi FM, Grattarola D, Alippi C. Spectral clustering with graph neural networks for graph pooling. In: International conference on machine learning. PMLR; 2020. p. 874–883.

[22] Wang X, Allen WE, Wright MA, Sylwestrak EL, Samusik N, Vesuna S, et al. Three-dimensional intact-tissue sequencing of single-cell transcriptional states. Science. 2018;361(6400):eaat5691.

[23] Uylings HB, Groenewegen HJ, Kolb B. Do rats have a prefrontal cortex? Behavioural brain research. 2003;146(1-2):3–17.

[24] Curtis C, Shah SP, Chin SF, Turashvili G, Rueda OM, Dunning MJ, et al. The genomic and transcriptomic architecture of 2,000 breast tumours reveals novel subgroups. Nature. 2012;486(7403):346–352.

[25] Birk S, Bonafonte-Pardàs I, Feriz AM, Boxall A, Agirre E, Memi F, et al. Quantitative characterization of cell niches in spatially resolved omics data. Nature Genetics. 2025; p. 1–13.

[26] Ma Y, Wang Y, Ma X. Signal-based spatial domain identification of spatially resolved transcriptomics with multigraph fusion. Briefings in Bioinformatics. 2026;27(1):bbag052.

