## Supplementary Materials for "Concordia: Spatial Domain Detection via Augmented Graphs for Population-Level Spatial Proteomics"

April 20, 2026

#### Supplementary Figures

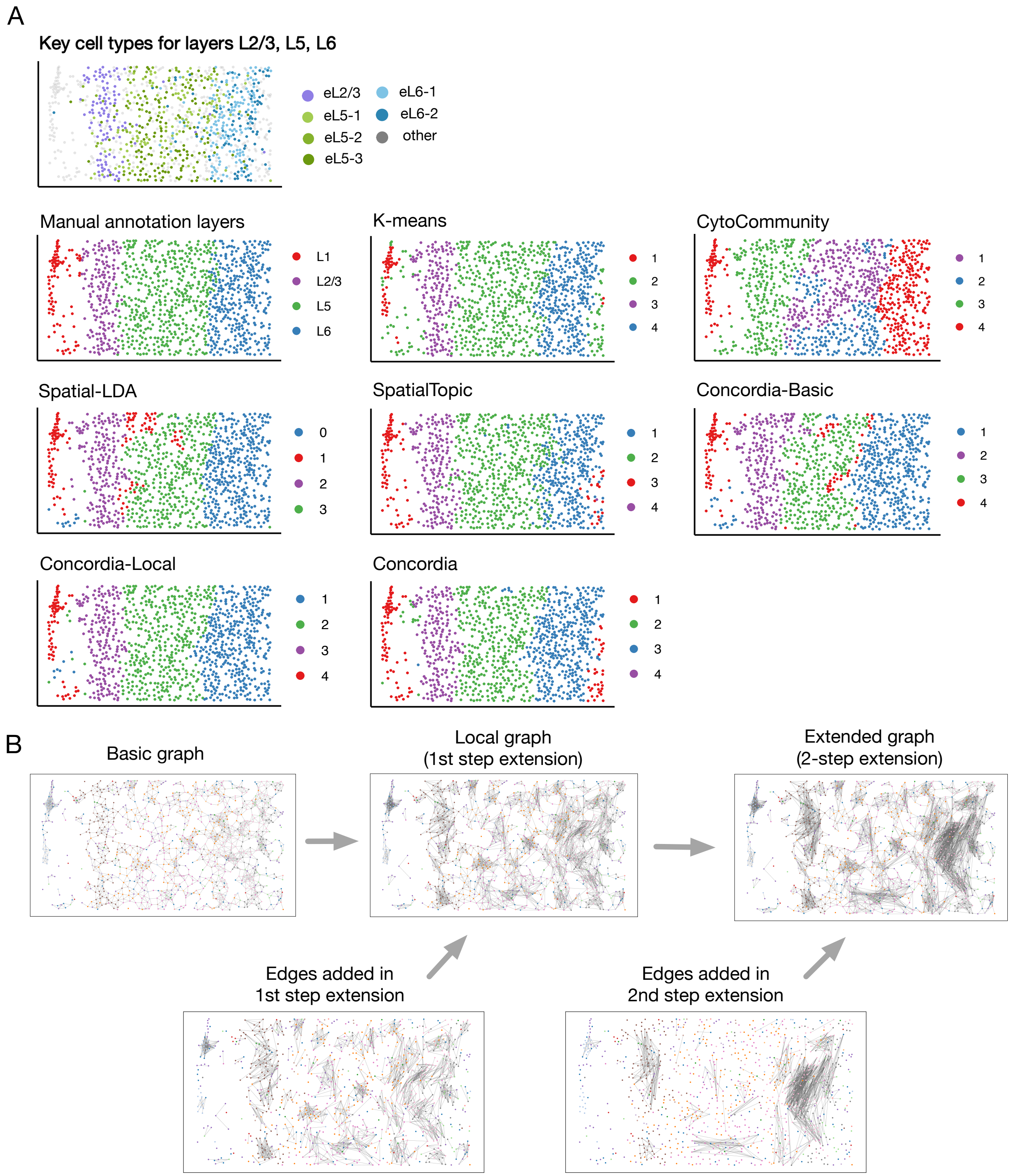


Supplementary Figure 1: Mouse medial prefrontal cortex tissue sample with ID 20180417_BZ5. (A) Distribution of key excitatory neuron cell types for layers L2/3, L5 and L6, manual layer annotation labels, and domain annotations from each method. (B) Three types of graphs and additional edges added in the graph extension steps.

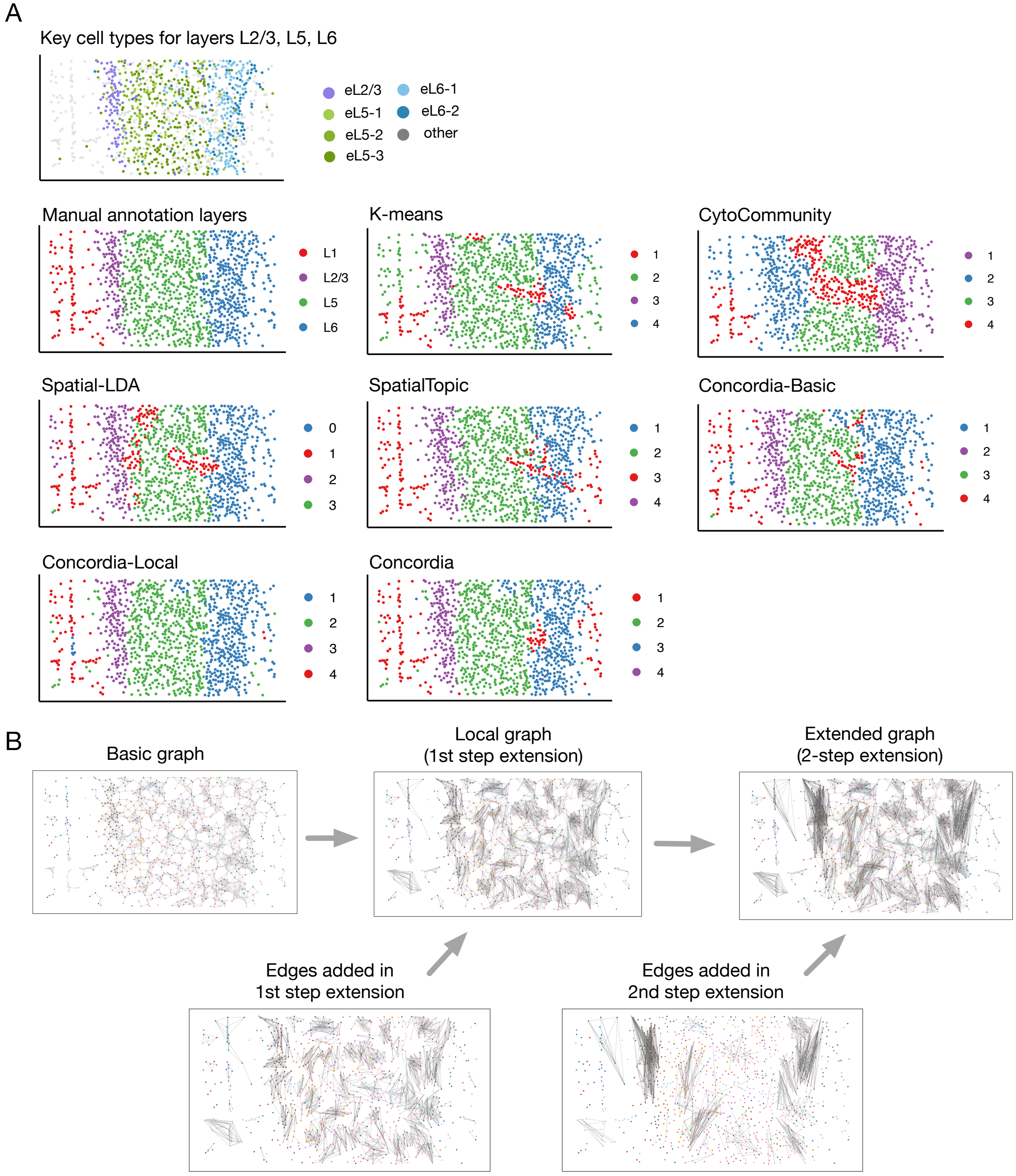


Supplementary Figure 2: Mouse medial prefrontal cortex tissue sample with ID 20180419_BZ9. (A) Distribution of key excitatory neuron cell types for layers L2/3, L5 and L6, manual layer annotation labels, and domain annotations from each method. (B) Three types of graphs and additional edges added in the graph extension steps.

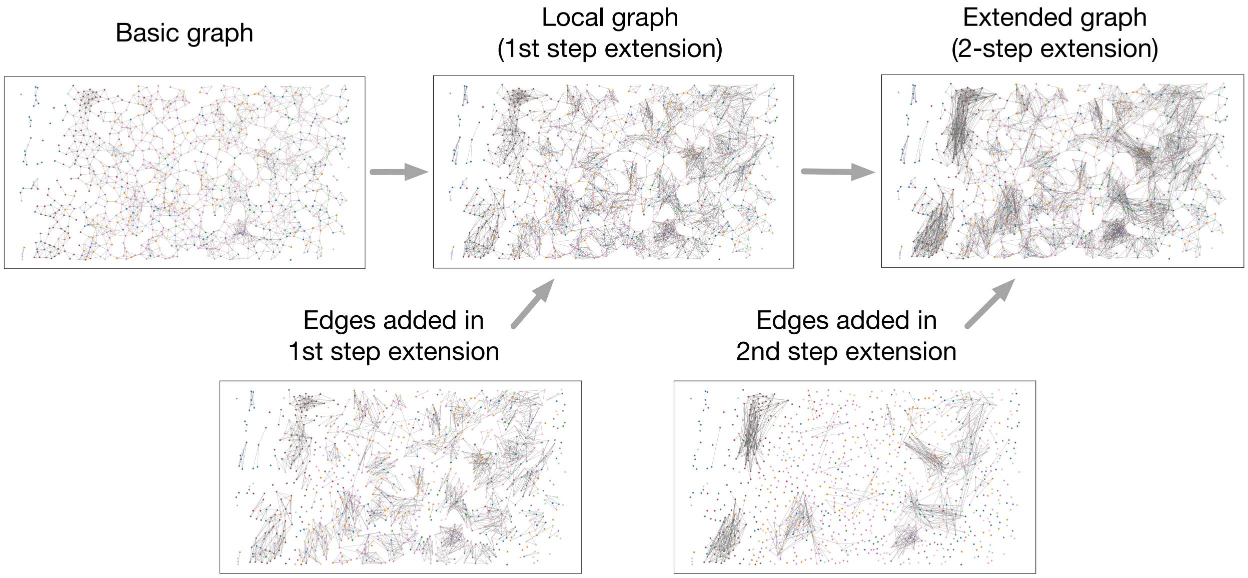


Supplementary Figure 3: Three types of graphs and additional edges added in the two steps of graph extension for mouse medial prefrontal cortex tissue sample with ID 20180424_BZ14.

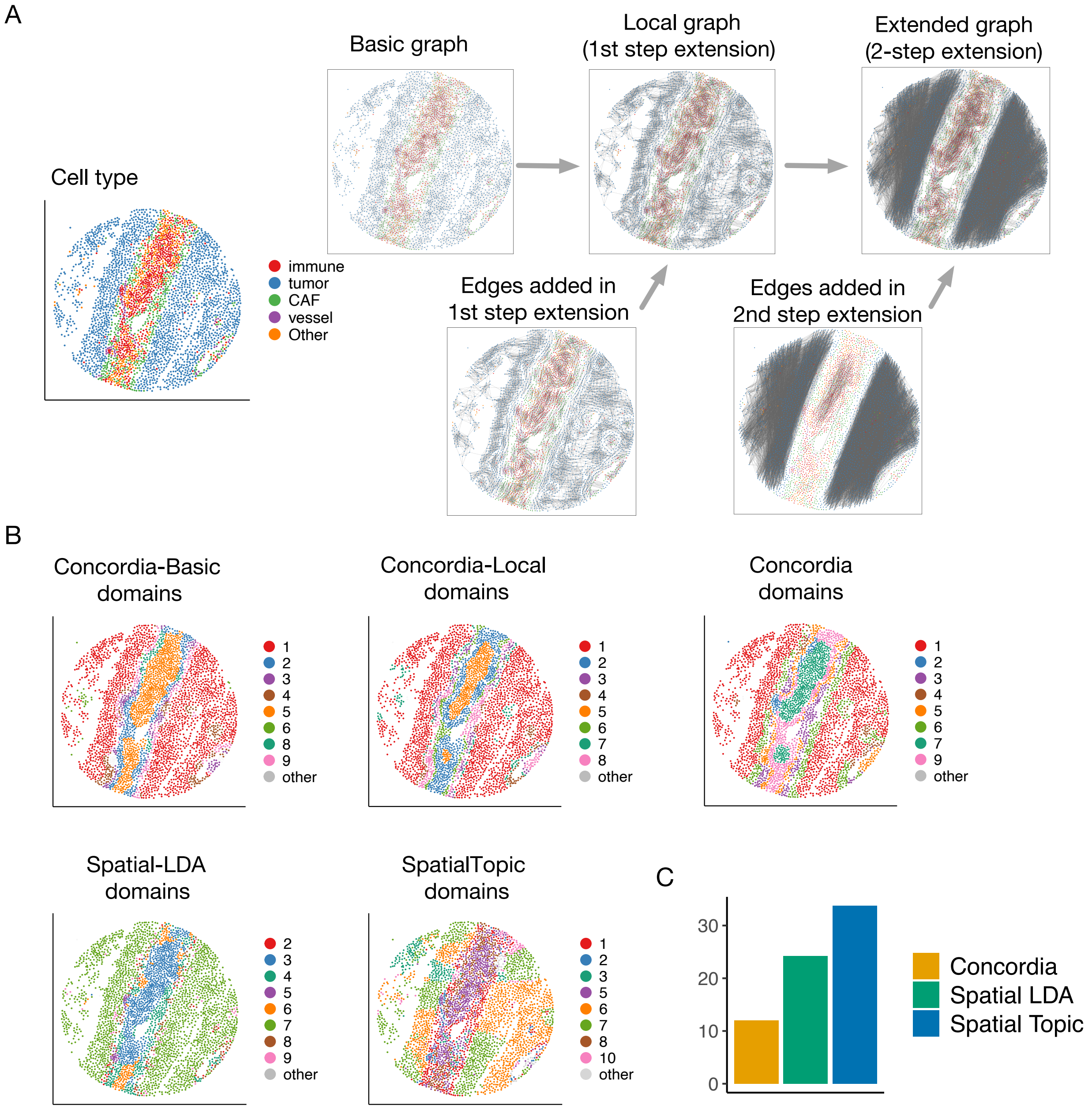


Supplementary Figure 4: Comparisons of different methods on an example image from lung cancer dataset. (A) Illustration of graph extension steps of Concordia on the example image, including three types of graphs. (B) Illustration of domain annotations given by five methods. Only domains that have at least 30 cells in the image are highlighted in each panel. Other domains are combined and labeled as “other”. (C) Barchart for the average number of connected components in domains based on the example image.

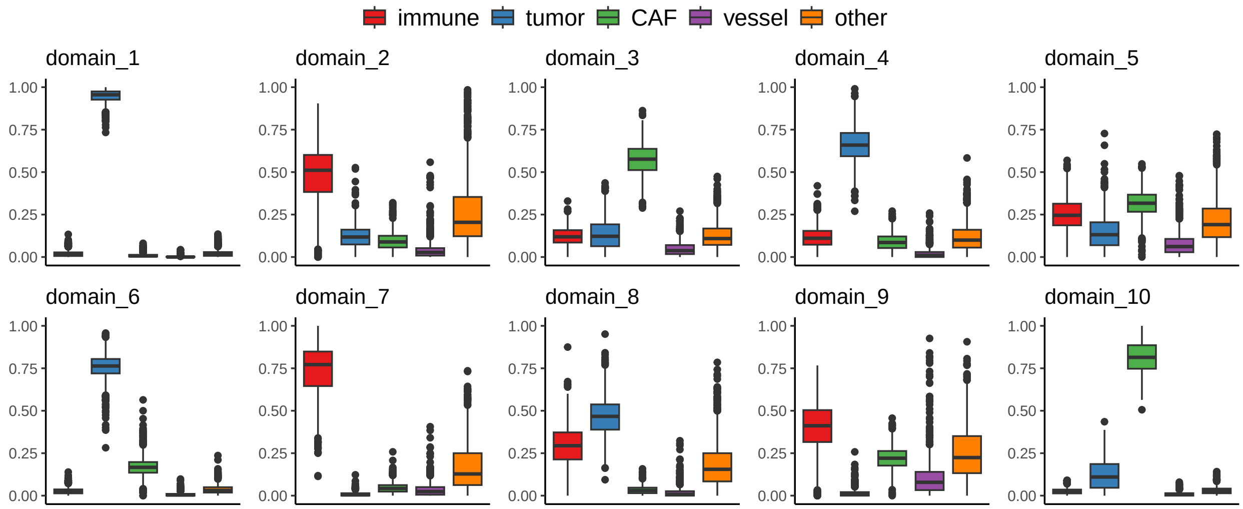


Supplementary Figure 5: Spatial domain characterization in lung cancer cohort. Boxplots of cell type composition within each of 10 domains identified by Concordia method (each point is one qualified image).

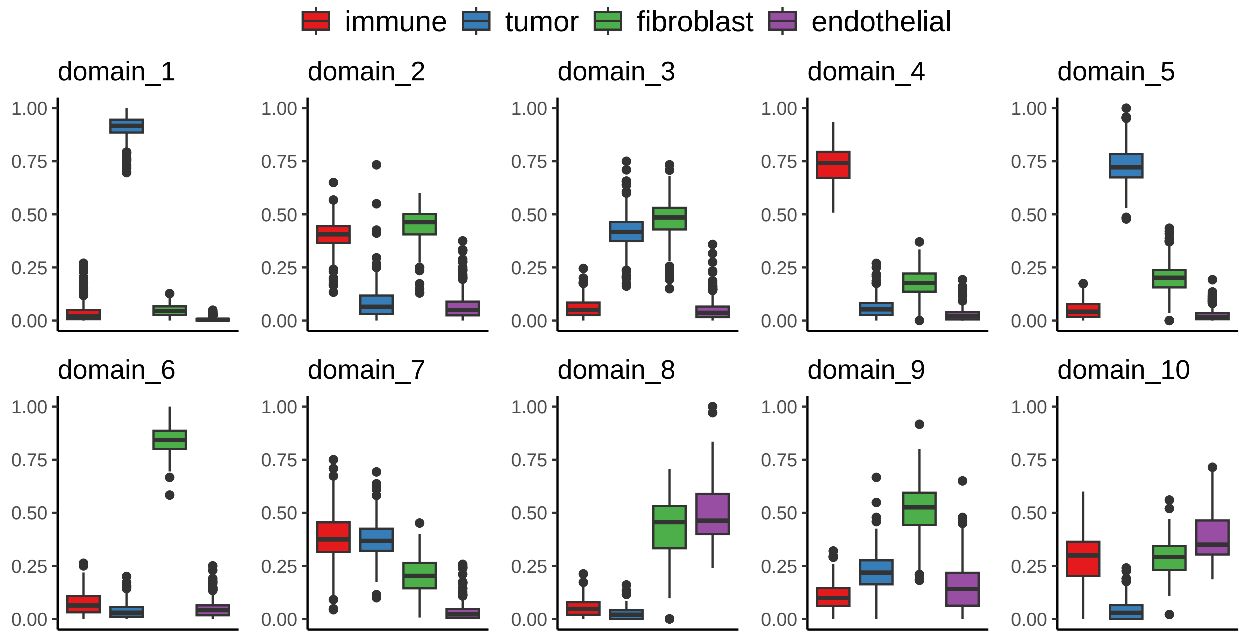


Supplementary Figure 6: Spatial domain characterization in breast cancer cohort. Boxplots of cell type composition within each of 10 domains identified by the Concordia method (each point is one qualified image).

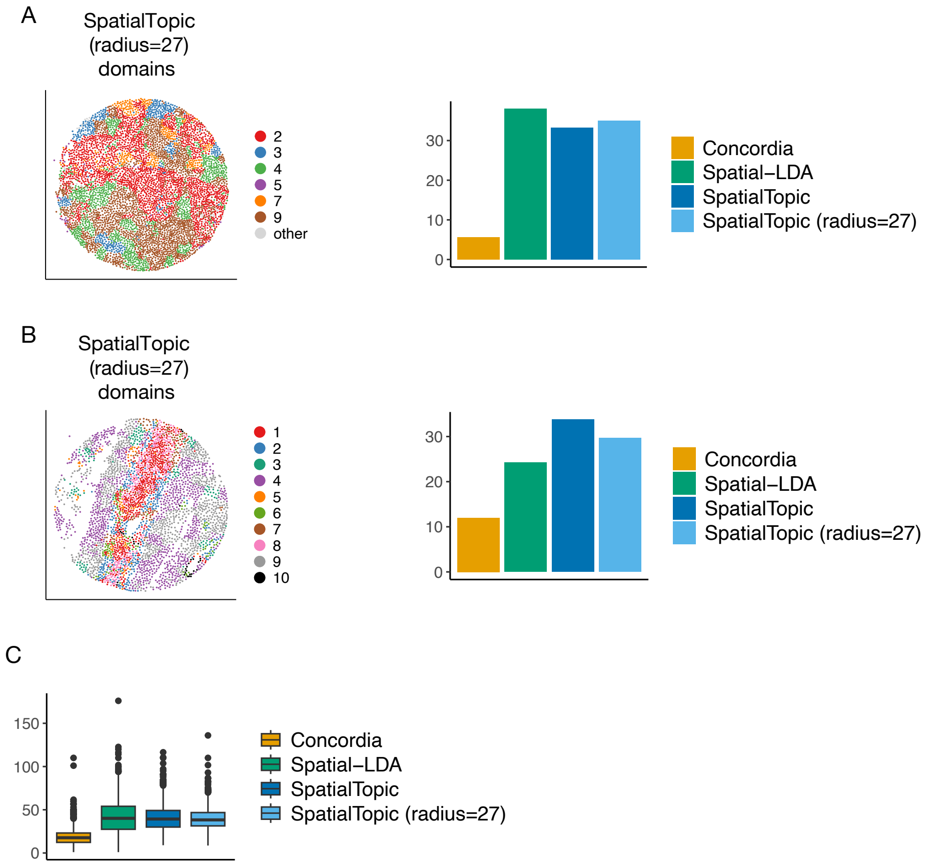


Supplementary Figure 7: On lung cancer dataset, illustration of results involving the domain annotation from SpatialTopic with radius=27. The radius value corresponds to each cell having average number of neighbors around 20 within the radius. (A) On the same image used in Fig. 3, domain annotation from SpatialTopic with radius=27 and barchart of the average number of connected components from the domains. Highlighted domains each has at least 30 cells. (Other domains are combined and labeled as "other". (B) On the same image used in Supplementary Fig. 4, domain annotation from SpatialTopic with radius=27 and barchart of the average number of connected components from the domains. (C) On all images from lung cancer dataset, boxplots for the average number of connected components given by four different methods including SpatialTopic with radius=27.

### Supplementary Table

Supplementary Table 1: ARI on images from Mouse brain dataset based on domain annotations given by different methods

| | 20180417_BZ5 | 20180419_BZ9 | 20180424_BZ14 |
|---|---|---|---|
| K-means | 0.653 | 0.520 | 0.670 |
| Concorida-Basic | 0.583 | 0.643 | 0.679 |
| Concorida-Local | 0.774 | 0.722 | 0.795 |
| Concordia | 0.671 | 0.669 | 0.720 |
| CytoCommunity | 0.399 | 0.354 | 0.456 |
| Spatial-LDA (radius_490) | 0.640 | 0.598 | 0.769 |
| Spatial-LDA (radius_690) | 0.700 | 0.631 | 0.811 |
| Spatial-LDA (radius_1600) | 0.516 | 0.474 | 0.682 |
| SpatialTopic (radius_490) | 0.765 | 0.659 | 0.697 |
| SpatialTopic (radius_690) | 0.804 | 0.710 | 0.822 |
| SpatialTopic (radius_1600) | 0.668 | 0.708 | 0.724 |

### Details of Methods

The details of the unsupervised clustering methods involved are listed in this section. Let $m \in \{1, \ldots, M\}$ represent the index of an image in a dataset, $i \in \{1, \ldots, N_m\}$ be the index of a cell among all $N_m$ cells in the $m$th image, and $c(i)$ represent the index of the cell type of cell $i$ among all $C$ cell types in the dataset.

**Basic graph** Build a graph for each image, with each cell as a node and two nodes are connected if their distance in coordinate space is no greater than the given distance cutoff. For each dataset, we randomly sampled a subset of images and selected the distance cutoff $r$ such that the median (across sampled images) of the resulting graph's average node degree was approximately 6. Let $d_{ij}$ be the distance in the coordinate space between cells $i$ and $j$ from the same image, given distance cutoff $r$, the basic graph is built by thresholding the distance between any two cells by $r$. For image $m$, the graph $G_0^{(m)} = (V^{(m)}, E_0^{(m)})$, where $V^{(m)} = \{1, \ldots, N_m\}$ and $E_0^{(m)} = \{\{i, j\} \mid d_{ij} \leq r, i \neq j, i \in \{1, \ldots, N_m\}, j \in \{1, \ldots, N_m\}\}$. All edges are undirectional.

In the basic graph, the set of direct neighbors of cell $i$ are all cells within $r$ distance, excluding $i$ itself. Using the notation of the graph edges, it can be written as

$$N_1^{(m)}(i) = \{j \in \{1, \ldots, N_m\} \mid \{i, j\} \in E_0^{(m)}, j \neq i\}.$$

The set of the second order neighbors of cell $i$ consists all cells that are first order neighbors of any cell in $N_1(i)$, excluding those in $N_1(i)$ and also cell $i$ itself. The set of the second order neighbors can be written as

$$N_2^{(m)}(i) = \{h \in \{1, \ldots, N_m\} \mid \{j, h\} \in E_0^{(m)} \; \forall j \in N_1(i), h \notin N_1(i), h \neq i\}.$$

The **2-hop neighborhood** is a concept involved in computing cell neighborhood similarity (i.e., neighborhood cell type composition or neighborhood CTC). For computing cell neighborhood similarity, the graph used is always the **basic graph** decided by distance cutoff in the coordinate space. For each cell, 2-hop neighborhood is the set of nodes containing the cell itself, the cells that are direct neighbors of the center cell, and those that are second order neighbors of the center cell. It can be written as

$$N_{\text{2-hop}}(i) = \{i\} \cup N_1(i) \cup N_2(i).$$

Between two cells in the same image, the cell neighborhood composition distance is computed based on the cell type composition in the 2-hop neighborhood. For each cell $i$, the cell type composition vector has length $C$, with the value at position $t$ being the proportion of cells in the 2-hop neighborhood that have cell type being the $t$th cell type, $t = \{1, \ldots, C\}$. In formula, for cell $i$, this vector is

$$c_i = \left(\frac{\sum_{h \in N_{\text{2-hop}}(i)} \mathbb{1}\{c(h) = 1\}}{|N_{\text{2-hop}}(i)|}, \frac{\sum_{h \in N_{\text{2-hop}}(i)} \mathbb{1}\{c(h) = 2\}}{|N_{\text{2-hop}}(i)|}, \ldots, \frac{\sum_{h \in N_{\text{2-hop}}(i)} \mathbb{1}\{c(h) = C\}}{|N_{\text{2-hop}}(i)|}\right).$$

The cell neighborhood distance between cell $i$ and cell $j$ is computed as L2 distance between the corresponding vectors, that is to say

$$comp\_dist(i, j) = \|c_i - c_j\|_2.$$

The smaller $comp\_dist(i, j)$ is, the similar cells $i$ and $j$ are in terms of the cell type composition in the 2-hop neighborhood.

**Local graph** (1st step extension only) Starting from the basic graph, for each node (cell), consider other cells within coordinate distance in the range $(r, 3 * r]$ from the node, add undirectional edges between the center cell and the top certain number of cells most similar to the center cell in terms of cell type composition in the 2-hop neighborhood (i.e., neighborhood cell type composition or neighborhood CTC). By default, the number of top cells is set to be 4. In addition, we put a threshold on the distance in cell type composition. In image $m$, For each cell $i$, write the set of cells within the required distance range as

$$candidates_{1st}^{(m)}(i) = \{h \mid d_{ih} \in (r, 3 * r], h \in \{1, \ldots, N_m\}\}.$$

Among all the cells in $candidates_{1st}^{(m)}(i)$, we rank their corresponding $comp\_dist(i, h)$ values in non-decreasing order. Let $comp\_dist\_4th$ be the 4th smallest distance value and $\epsilon$ be the additional cutoff on distance in cell type composition, then for cell $i$, the additional edges to add are

$$E_{1stadd}^{(m)}(i) = \{\{i, h\} \mid h \in candidates_{1st}^{(m)}(i), comp\_dist(i, h) \le min(comp\_dist\_4th, \epsilon)\}.$$

For the $m$th image, all additional edges to add are

$$E_{1stadd}^{(m)} = \cup_{i \in \{1, \ldots, N_m\}} E_{1stadd}^{(m)}(i).$$

The graph with step 1 extension only is $G_{1st}^{(m)} = (V^{(m)}, E_{1st}^{(m)})$, where

$$E_{1st}^{(m)} = E_0^{(m)} \cup E_{1stadd}^{(m)}.$$

By default, the 2-hop neighborhood cell type composition distance threshold $\epsilon$ is set to be 0.176, slightly lower than the value of $\sqrt{2 \cdot (1/8)^2}$. $\sqrt{2 \cdot (1/8)^2}$ corresponds to the case when 1/8 of cells in 2-hop neighborhoods of cell $i$ are in one cell type and 1/8 of cells in that of cell $h$ are in a different cell type, while the values in $c_i$ and $c_h$ on positions for other cell types are the same.

**Extended graph** (after two steps of extension) Starting from the graph with step 1 extension only, further augment the graph by adding more qualified edges. The number of edges to add is chosen so that the average degree of nodes in each graph (image) meets a pre-specified number. By default, the average degree 20 is used. An edge between two cells is qualified to be a candidate edge to add if: (1) the two cells at the ends are similar enough in terms of their 2-hop neighborhoods; (2) on the shortest path between these two cells, at least certain proportion of the cells on the path are similar enough to the two cells at the ends in terms of their 2-hop neighborhoods. By default, the proportion threshold is chosen to be 90%. Here, the shortest path is computed based on the edges in the graph after the step 1 extension, i.e., $G_{1st}^{(m)} = (V^{(m)}, E_{1st}^{(m)})$, and the similarity in terms of the 2-hop neighborhoods is reflected by the L2 distance between the cell type composition vectors of the two cells' 2-hop neighborhoods in the basic graph, i.e., $comp\_dist(i, j)$ for cell $i$ and cell $j$, and $c_i$ and $c_j$ are computed based on the 2-hop neighborhoods in the basic graph $G_0^{(m)}$ for each cell. Cells $i$ and $j$ are viewed as similar enough in terms of their 2-hop neighborhoods if $comp\_dist(i, j)$ is below a specified threshold. By default, this threshold is the same as the $\epsilon$ used in the 1st step extension only graph.

In one graph (image), there can be more edges qualified as candidates to add to the graph than the number of edges needed to meet the desired average degree. We sample the desired number of additional edges from the qualified ones. To encourage more long-distance connections, we do stratified sampling by binning the candidate edges into groups according to the path length and sampling more edges from the long-distance groups to add to the graph. In detail, we bin all candidate edges into four groups separated by quartiles of their path lengths. Among the total number of additional edges needed to meet the specified average degree, we get 10% of them by sampling from the bin of path length no higher than 25 percentile, 20% from the bin of path length between 25% percentile and the median, 30% from the bin of path length between the median and 75 percentile, and 40% from the bin of path length above 75 percentile. If for any bin, there are not enough number of candidate edges to sample from it, we include all edges from this bin. The resulting graph is the fully extended graph, and we write it as $G_{extended}^{(m)} = (V^{(m)}, E_{extended}^{(m)})$.

**Input node feature to the GNN model** The feature vector for each cell is concatenation of three vectors, each representing the cell type composition based on the center cell itself, the direct neighbors, and the second order neighbors. The vector for the center cell itself is a one-hot encoded vector, with 1 only for the position corresponding to the cell type of the center cell and 0 everywhere else. The neighborhood for computing the components of feature vector changes according to the model setting (basic graph, local graph, or extended graph). In each setting, the same graph is used for obtaining input node feature, graph convolution layers, and unsupervised loss function.

**Combine small clusters into large domains** To combine the small clusters obtained from running $K$-means on embedding space, we consider two types of distances between any two clusters, the distance in embedding space and the distance in physical space. Given any two clusters, for both types of distances, the distances are summarized only based on the images that have enough number of cells for both of the clusters. A distance matrix is formed for each of the two distance types between all small clusters. The final weighted distance matrix is a weighted average, where the weight is used to adjust for the scale difference between the distances from the two aspects. The final domains are decided by hierarchical clustering among all clusters based on the weighted distance matrix. According to specified desired number of domains, the hierarchical clustering gives the mapping from small clusters to large domains. For example, if a domain corresponds to five clusters, then all cells under these five clusters are considered to be in the given domain.

In formula, let $a$ and $b$ represent two $K$-means clusters, and let $a^{(m)}$ and $b^{(m)}$ represent the cells that they occupy in image $m$, respectively. Under a given cluster size cutoff $n_{min}$, in image $m$, if $a^{(m)}$ and $b^{(m)}$ each occupies at least $n_{min}$ cells, we say that they co-occur in image $m$. For each dataset, $n_{min}$ was selected based on the typical number of cells per image to account for differences in image scale from different datasets. Write the set of images in which the clusters $a$ and $b$ co-occur as $cooccur_{ab} = \{m \in \{1, \ldots, M\} \mid |a^{(m)}| \geq n_{min}, |b^{(m)}| \geq n_{min}\}$.

To ensure that the between-cluster distances computed are robust, we only consider the pairs of clusters co-occuring in enough number of images. In each dataset, the number of images cutoff $N_{img}$ was selected based on the total number of images to account for differences in cohort scale from different dataset. Under the specified number of images cutoff $N_{img}$, if clusters $a$ and $b$ co-occur in at least $N_{img}$ images, i.e., $|cooccur_{ab}| \geq N_{img}$, we proceed to compute the embedding and physical distances between these two clusters. First, we compute distances in each image from the set $cooccur_{ab}$. In an image with index $m \in cooccur_{pq}$, the embedding distance between clusters $a$ and $b$ is computed as the L2 distance in the embedding space between the centroids of embedding features of the two sets of cells:

$$d_{emb}^{(m)}(a, b) = \|\mu_a^{(m)} - \mu_b^{(m)}\|_2,$$

where $\mu_a^{(m)}$ and $\mu_b^{(m)}$ are embedding centroids of clusters $a$ and $b$ in image $m$, respectively.

On the other hand, in image $m$, to compute physical distance, we first get the directional shortest distances in the coordinate space between the cells in $a^{(m)}$ and those in $b^{(m)}$. In details, without loss of generality, assuming the indices of the cells in $a^{(m)}$ are $1, 2, \ldots, |a^{(m)}|$, then the vector of shortest distances from the cells in $a^{(m)}$ to the cells in $b^{(m)}$ is given by

$$d_{coord\_a \to b}^{(m)} = (\delta(1, b^{(m)}), \delta(2, b^{(m)}), \ldots, \delta(|a^{(m)}|, b^{(m)})),$$

where $\delta(i, b^{(m)})$ is the Euclidean distance from $i$ to its nearest cell in cluster $b$ in image $m$, i.e., $\delta(i, b^{(m)}) = \min_{j \in b^{(m)}} d_{ij}$, and $d_{ij}$ is the L2 distance in the coordinate space between cell $i$ and cell $j$. The vector of shortest distances from the cells in $b^{(m)}$ to the cells in $a^{(m)}$ is computed similarly. Then, write the 90% quantile values of $d_{coord\_a \to b}^{(m)}$ and $d_{coord\_b \to a}^{(m)}$ as $q_{a \to b}^{(m)}$ and $q_{b \to a}^{(m)}$, respectively, and the physical distance between clusters $a$ and $b$ in image $m$ is the minimum of these two values:

$$d_{\text{coord}}^{(m)}(a, b) = \min\left(q_{a \to b}^{(m)}, q_{b \to a}^{(m)}\right),$$

The embedding distance $d_{emb}(a, b)$ and physical distance $d_{coord}(a, b)$ at the data set level between clusters $a$ and $b$ are the median of the corresponding distance values in all images that these two clusters co-occur in (with enough number of cells), respectively. This is to say:

$$d_{emb}(a,b) = \text{median}_{m \in cooccur_{ab}} d^{(m)}_{\text{emb}}(a,b)$$

$$d_{coord}(a,b) = \text{median}_{m \in cooccur_{ab}} d^{(m)}_{\text{coord}}(a,b).$$

Next, we compute a distance matrix between all pairs of clusters. This distance matrix $\mathbf{D}_{K\times K}$, where $K$ is the total number of $K$-means clusters, combines the embedding distance and physical distance. Due to that these two types of distances can be on different scales, we use a weight factor to bring the median embedding distance across all qualified pairs of clusters to be the same as the median coordinate distance, and then take an average. For each pair of clusters $a$ and $b$ co-occuring in enough number of images, i.e., $|cooccur_{ab}| >= N_{img}$, the final distance value, $\mathbf{D}_{ab}$, which is an element in the matrix $\mathbf{D}$, is computed as:

$$\mathbf{D}_{ab} = (w * d_{emb}(a,b) + d_{coord}(a,b))/2,$$

where weight factor $w$ is decided by the ratio of the median embedding distance and the median physical distance across all qualified cluster pairs. This is to say, if we write the set of qualified cluster pairs as:

$$P = \{(a,b) \mid |cooccur_{ab}| >= N_{img}, a \in \{1,2,\dots,K\}, b \in \{1,2,\dots,K\}, a \neq b\},$$

then,

$$w = \frac{median_{(a,b)\in P}\ d_{coord}(a,b)}{median_{(a,b)\in P}\ d_{emb}(a,b)}.$$

For any cluster pair $(a,b) \notin P$, where $a \neq b$, they do not co-occur in enough number of images, and thus they do not have defined embedding distance and physical distance. In this case, we do a linear interpolation between $\max(\{\mathbf{D}_{st} \mid (s,t) \in P\})$ and $2 * \max(\{\mathbf{D}_{st} \mid (s,t) \in P\})$ according to the number of images they co-occur in, with the interpolated distance being $2 * \max(\{\mathbf{D}_{st} \mid (s,t) \in P\}$ when they do not co-occur in any image:

$$\mathbf{D}_{ab} = \max(\{D_{st} \mid (s,t) \in P\}) * \frac{2 * N_{img} - |cooccur_{ab}|}{N_{img}}.$$

All diagonal elements $\mathbf{D}_{aa}, a \in \{1,2,\dots,K\}$, are set to 0 since they each corresponds to the pair between one cluster and itself.

To obtain final domains, hierarchical clustering on the $K$-means clusters is done based on the distance matrix $\mathbf{D}$ with the complete linkage method (`hclust` function in `R` with `method="complete"`). At a specified number of domains, the dendrogram is cut to obtain the correspondence between the $K$-means clusters and the resulting domains.

**Dataset-specific thresholds.** Three thresholds were set in a dataset-specific manner, which are the radius $r$ for constructing the base distance-threshold graph and two eligibility criteria for computing inter-cluster distances: two clusters need to co-occurs in at least $N_{img}$ images and, in each of those images, both clusters contain at least $n_{min}$ cells. For lung cancer dataset, $r = 16$, $n_{min} = 30$ and $N_{img} = 30$. For breast cancer dataset, $r = 20$ and $n_{min} = 20$ and $N_{img} = 10$. For mouse medial prefrontal cortex dataset, $r = 380$, $n_{min} = 10$ and $N_{img} = 1$.
